# *Listeria* spp. Isolated from Soil Samples Collected in the Great Smoky Mountains

**DOI:** 10.1101/2021.11.19.469259

**Authors:** Michelle L. Claxton, Lauren K. Hudson, Daniel W. Bryan, Thomas G. Denes

## Abstract

*Listeria monocytogenes,* a foodborne pathogen, and other *Listeria* spp. are present in natural environments. Isolating and characterizing strains from natural reservoirs can provide insight into the prevalence and diversity of *Listeria* spp. in these environments, elucidate their contribution to contamination of agricultural and food processing environments and food products, and lead to the discovery of novel species. In this study, we evaluated the diversity of *Listeria* spp. isolated from soil samples in a small region of the Great Smoky Mountains National Park (GSMNP), which is the most biodiverse national park in the United States National Park system. Of the 17 *Listeria* isolates that were recovered, whole-genome sequencing revealed that 14 were unique strains. The unique strains were shown to represent a diversity of *Listeria* spp., including *L. monocytogenes* (n=9), *L. cossartiae* subsp. *cossartiae* (n=1), *L. marthii* (n=1), *L. booriae* (n=1), and a novel Listeria sp. (n=2). The *Listeria* isolated in this study were collected from high elevation sites near a creek that drains into a series of rivers ultimately leading to the Mississippi River; thus, the *Listeria* present in this natural environment could potentially travel downstream to a large region that includes portions of nine southeastern and midwestern states in the U.S. The *Listeria* spp. isolated and described in this study provide insight into the diversity of *Listeria* spp. found in the Great Smoky Mountains and indicate that this environment is a reservoir of novel *Listeria* spp.

**IMPORTANCE:** *Listeria monocytogenes* is a foodborne pathogen that can cause serious systemic illness that, although rare, usually results in hospitalization and has a relatively high mortality rate compared to other foodborne pathogens. Identification of novel and diverse *Listeria* spp. provides insight into the genomic evolution, ecology, and evolution and variance of pathogenicity of this genus, especially in natural environments. Comparing *L. monocytogenes* and *Listeria* spp. isolates from natural environments, such as those recovered in this study, to contamination and/or outbreak strains may provide more information about the original natural sources of these strains and the pathways and mechanisms that lead to contamination of food products and agricultural or food processing environments.

## INTRODUCTION

Members of the genus *Listeria* are characterized as Gram-positive, short rods, motile via peritrichous flagella, and aerobic or facultative anaerobic metabolisms (1). The *Listeria* genus currently contains 26 validly published species (as of October 2021) (2, 3). Of these, only two are typically described as pathogenic to humans: *L. monocytogenes*, a human and animal pathogen, and *L. ivanovii*, which primarily infects ruminants, but is an extremely rare opportunistic human pathogen (1, 4-9). Two other species, *L. seeligeri* and *L. innocua*, have each been documented to cause disease in humans only once (10, 11). Listeriosis in humans is primarily caused by ingesting food contaminated with *L. monocytogenes* (1). Compared to other foodborne pathogens, the incidence of listeriosis is low (0.3 cases per 100,000 population); however, the proportion of cases resulting in hospitalization (98%) or death (16%) are high (12), as are the economic costs associated with this pathogen (estimated as up to $8.6 billion) (13).

As *L. monocytogenes* is a foodborne pathogen, much research has focused on evaluating the distribution of *L. monocytogenes* and other *Listeria* spp. in food, food-processing facilities, agricultural settings, and animals. However, *Listeria* spp. are distributed in a variety of natural environments, including soil, sewage, animal feces, agricultural environments, decaying vegetation, and water (1, 14-17). The adaptability of this microorganism plays a role in its wide distribution: *Listeria* spp. can survive in a wide range of pH (4.5-9.2), temperature (0-45 C°), and salt concentrations (≤10% NaCl) (18). The extensive variety of natural reservoirs cause variable genomic flexibility and recombination (19). There is interest in describing the distribution, prevalence, variability, diversity, and characteristics of *Listeria* spp. from natural environments (17). A better understanding of the ecology and evolution of *Listeria* spp. will help to elucidate the sources and pathways that lead to contamination of agricultural products or food processing facilities (20). Additionally, sampling in agricultural and natural environments can lead to the discovery of novel *Listeria* spp. Since 2010, 12 novel *Listeria* spp. isolated from natural environments have been described (14, 21-24). This indicates that novel species of *Listeria* strains can be found by surveying biodiverse natural reservoirs.

A recent study examined *Listeria* spp. in soil across the contiguous United States and reported a 31% prevalence of Listeria spp. in the soil samples collected and that the most prevalent *Listeria* phylogroup was L8 (*L. welshimeri*), followed by L1(*L. monocytogenes* lineage III) and L12 (*L. booriae*) (*L. monocytogenes* would be the most prevalent if all three lineage phylogroups are considered as a whole) (15). The geographical distance between strains has also been shown to correlate with the gene content difference, but only minimally in *Listeria* spp. (15, 19). In a study conducted in Australia, *L. seeligeri, L. innocua*, and *L. ivanovii* were the most dominant species isolated from soil and water samples (25).

The purpose of this study was to evaluate the diversity of *Listeria* spp. isolated from soil samples in a small region of the Great Smoky Mountains, a subrange of the Appalachian Mountains that is located along the Tennessee-North Carolina border and mostly contained within the Great Smoky Mountains National Park (GSMNP). A location within the GSMNP was chosen because of the park’s high level of biodiversity: it is the most biodiverse of all the parks in the United States National Park System and contains 19,000 documented species of animals, plants, fungi, and other organisms within the 800 square mile area, with an estimated additional 80,000-100,000 undocumented species (26). Other researchers have found that *Listeria* spp. prevalence is highest throughout and near the Mississippi River Basin (15). Additionally, the specific sampling location along the Twentymile Creek is of interest, as the downstream flow of includes prominent water systems, including the Tennessee and Mississippi Rivers, and runs through nine southeastern and midwestern states (27, 28). *Listeria* spp. present in this location may potentially be carried downstream to large areas of the region that contain people, livestock, farms, and food processing facilities.

## RESULTS AND DISCUSSION

Twelve soil samples were collected from twelve separate sites along the Twentymile Trail in GSMNP (**Figure 1**). A total of 17 *Listeria* spp. isolates were recovered from nine of the samples (**Table 1**). Of the 17 isolates, 14 were unique strains (those that were zero hqSNPs apart were considered identical). The most prevalent species was *L. monocytogenes* (n=10), with nine unique strains (UTK C1-0007 and UTK C1-0012 were identical) recovered from six different sites. Sites K and L are within fifty feet of one another, and both sites showed a prevalence of *L. monocytogenes*. Other species that were isolated included *L. cossartiae* subsp. *cossartiae* (n=2; one unique strain: UTK C1-0002 and UTK C1-0005 were identical), *L. marthii* (n=1), and *L. booriae* (n=1). All of these isolates had ANI values >96% with their respective type strains, which is above the 95% cut-off typically used for bacterial species distinction (29, 30). Three of the isolates (two unique strains; UTK C1-0017 and UTK C1-0024 were identical) clustered together but shared <95% ANI with all known *Listeria* spp. type strains, suggesting they would qualify as a new species.

**Figure 1.**
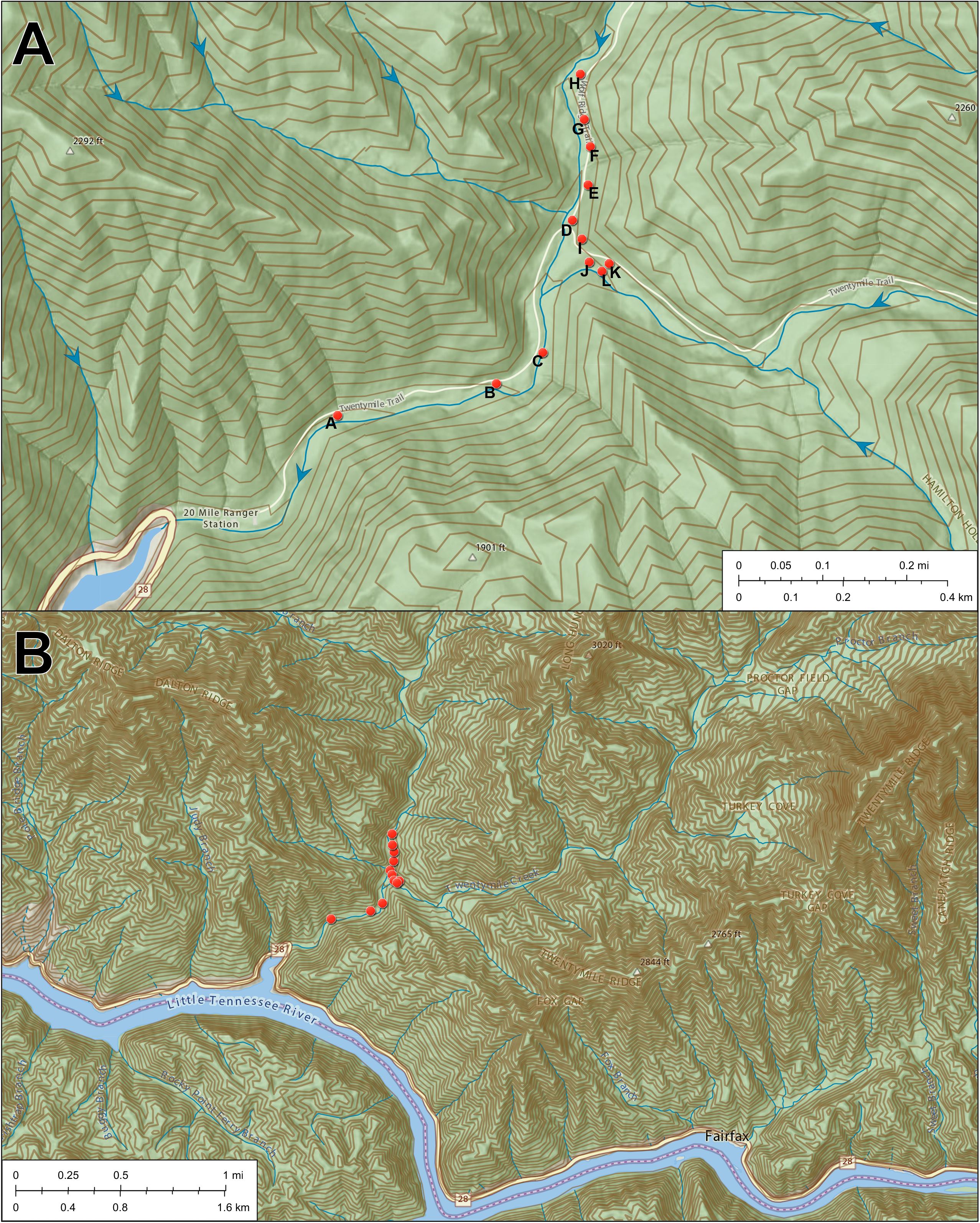
Maps of sampling location and features. Maps of sampling location and surrounding area with: (A) the Twentymile Loop trail with sampling sites (A-L) indicated with red circles and water flow direction indicated with blue arrows; and (B) the surrounding area around the sampling location. Topographical contours are indicated by brown lines, with higher elevations are indicated by thicker lines. The altitude ranges from 1,280 to 2,850 ft above sea level.

**Figure 2.**
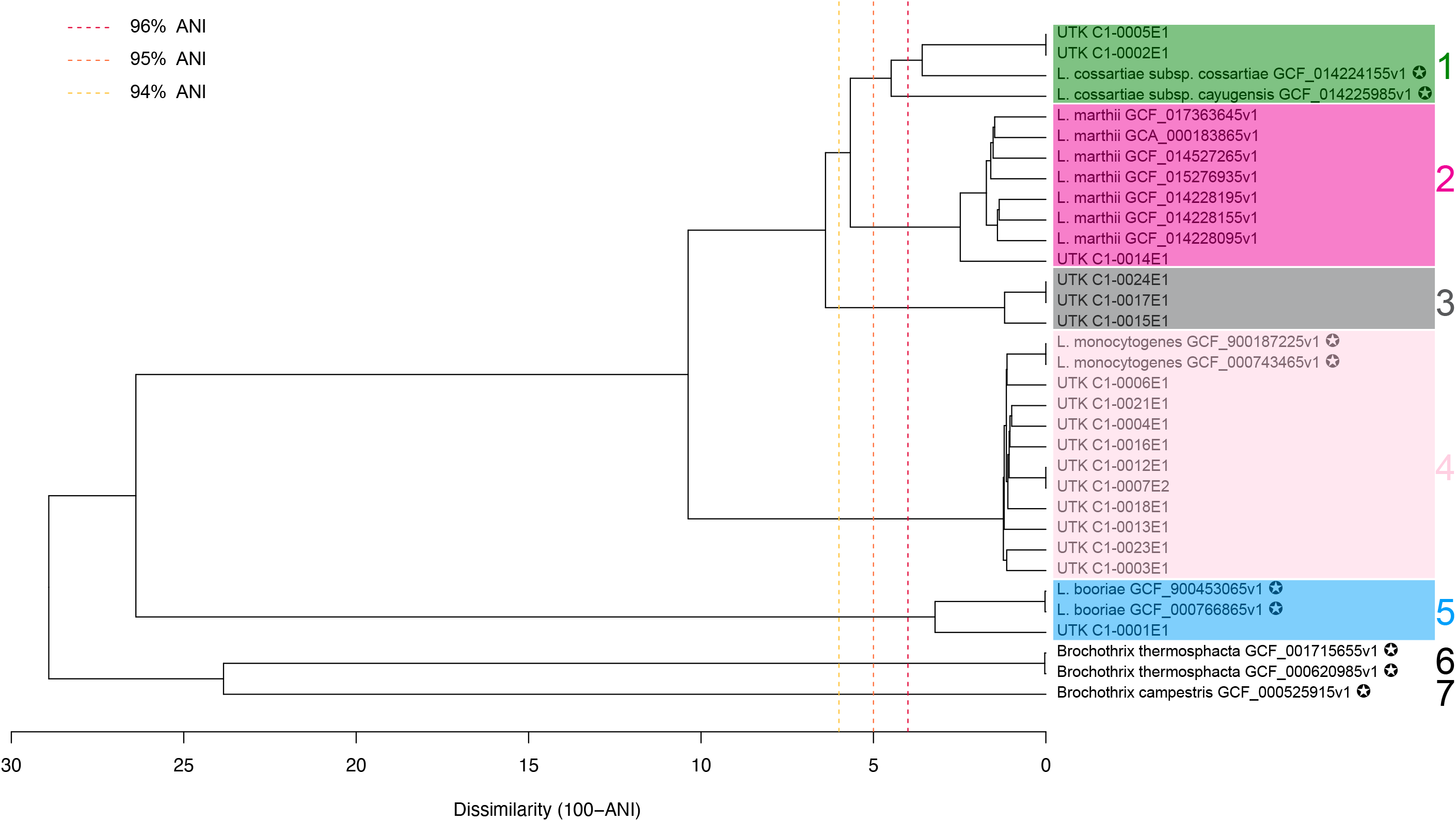
Average nucleotide identity (ANI) dendrogram. Dendrogram includes isolates from this study and type strains (indicated by a star). The dashed vertical lines indicate reference ANI values (see legend). Three *Brochothrix* spp. type strains were also included as an outgroup.

**Table 1.** *Listeria* spp. isolates recovered from soil samples collected in the Great Smoky Mountains National Park.

| Isolate | Species | Sample Site | Assembly Statistics |  |  |  | BioSample ID |
| --- | --- | --- | --- | --- | --- | --- | --- |
|  |  |  | Length (Mb) | G+C Content (%) | No. Contigs | Avg. Coverage (×) |  |
| UTK C1-0001 | *L. booriae* | A | 3.78 | 41.78 | 23 | 108.0 | SAMN21217721 |
| UTK C1-0002<sup>a</sup> | *L. cossartiae*<br>subsp. *cossartiae* | B | 2.86 | 38.72 | 15 | 154.2 | SAMN21217722 |
| UTK C1-0003 | *L. monocytogenes* | C | 2.99 | 37.67 | 19 | 134.5 | SAMN21217723 |
| UTK C1-0004 | *L. monocytogenes* | A | 3.04 | 37.66 | 13 | 154.1 | SAMN21217724 |
| UTK C1-0005<sup>a</sup> | *L. cossartiae*<br>subsp. *cossartiae* | B | 2.86 | 38.72 | 15 | 149.3 | SAMN21217725 |
| UTK C1-0006 | *L. monocytogenes* | C | 2.96 | 37.84 | 14 | 159.4 | SAMN21217726 |
| UTK C1-0007<sup>b</sup> | *L. monocytogenes* | F | 3.01 | 37.77 | 22 | 135.3 | SAMN21217727 |
| UTK C1-0012<sup>b</sup> | *L. monocytogenes* | F | 3.01 | 37.77 | 21 | 149.0 | SAMN21217728 |
| UTK C1-0013 | *L. monocytogenes* | L | 3.00 | 37.73 | 20 | 162.6 | SAMN21217729 |
| UTK C1-0014 | *L. marthii* | H | 2.83 | 38.52 | 12 | 180.8 | SAMN21217730 |
| UTK C1-0015 | *Listeria* spp. | I | 2.81 | 38.62 | 10 | 154.5 | SAMN21217731 |
| UTK C1-0016 | *L. monocytogenes* | H | 2.97 | 37.81 | 15 | 139.6 | SAMN21217732 |
| UTK C1-0017<sup>c</sup> | *Listeria* spp. | J | 2.92 | 38.5 | 11 | 153.6 | SAMN21217733 |
| UTK C1-0018 | *L. monocytogenes* | K | 3.15 | 37.64 | 19 | 73.4 | SAMN21217734 |
| UTK C1-0021 | *L. monocytogenes* | L | 3.04 | 37.69 | 18 | 141.3 | SAMN21217735 |
| UTK C1-0023 | *L. monocytogenes* | K | 2.92 | 37.76 | 17 | 145.5 | SAMN21217736 |
| UTK C1-0024<sup>c</sup> | *Listeria* spp. | J | 2.92 | 38.5 | 11 | 164.8 | SAMN21217737 |

<sup>a</sup>isolates are identical (hqSNP distance of 0)
<sup>b</sup>isolates are identical (hqSNP distance of 0)
<sup>c</sup>isolates are identical (hqSNP distance of 0)

The species identified in the present study correspond to four of the 12 phylogroups described by Liao, et al. (15): L2 (*L. monocytogenes* lineage II), L4 (*L. marthii*), L5 (*L. cossartiae*), and L12 (*L. booriae*). In that previous study, phylogroups L2 and L12 had a wide distribution across the United States, with L12 being isolated from the GSMNP region and L2 not. In contrast, phylogroups L4 and L5 had much smaller distributions, with L4 only being isolated in Kentucky, Pennsylvania and New York, and L5 only in Alabama (neither was isolated in the GSMNP region). *L. marthii* was originally isolated from soil and water samples collected in the Finger Lakes National Forest in New York (21) and *L. cossartiae* subsp. *cossartiae* was originally isolated from soil samples collected in Alabama (24). L1, L3, and L6 have previously been isolated in the GSMNP region (15), but were not isolated in the present study.

*L. monocytogenes* and other *Listeria spp*. exist in natural environments, including soil and on plants, and in the intestinal tracts of animals and can be spread to the environment via feces. Humans and animals within these environments are exposed to *Listeria* frequently (31). Produce has been implicated in several listeriosis outbreaks (32-38). Preharvest contamination of produce or agricultural environments can occur if *L. monocytogenes* is present in the soil or introduced via wildlife, raw or improperly composted manure, or irrigation water (38-40). All sample sites that tested positive for *L. monocytogenes* (sites A, C, F, H, K, and L) in the current study were within fifty feet of the Twentymile Creek. The downstream flow of this creek includes the Little Tennessee, Tennessee, Ohio, and Mississippi Rivers. Ultimately, the flow from Twentymile Creek to the Gulf of Mexico goes through nine states (North Carolina, Tennessee, Alabama, Kentucky, Illinois, Missouri, Arkansas, Mississippi, and Louisiana) (27, 28). This is significant because *L. monocytogenes* from the sampling location could enter the creek, be carried downstream, and potentially migrate to local or regional agricultural environments or irrigation water and contaminate produce or infect livestock, which could result in human illnesses or outbreaks.

Additionally, food processing environments can become contaminated with *Listeria* spp. which can lead to contamination of food products during processing. For the purpose of environmental monitoring programs in food processing facilities, testing for *Listeria* spp. is typically conducted as an indicator for the potential presence of *L. monocytogenes* and/or for monitoring the effectiveness of sanitation practices (41-44). Once established in the food processing environment, *Listeria* spp. may be able to persist for years (41, 45). *Listeria* spp. can be initially introduced into a food processing facility via food products, ingredients, equipment, humans, etc. (33, 46), but often, the original source of the strain is unknown.

As more *L. monocytogenes* and *Listeria* spp. isolates from natural environments are sequenced and characterized, such as was done in this and similar studies (15-17, 25), they can be compared with contamination and/or outbreak strains. Doing so may provide more information about the original natural sources of these isolates and the pathways and mechanisms that lead to contamination of food products and agricultural or food processing environments. For future studies, a larger and broader geographical region should be investigated to determine ecological and evolutionary characteristics of *Listeria* spp. Natural environments closer to agricultural locations should be investigated as well.

## MATERIALS AND METHODS

### Soil sample collection

Soil samples were collected along the Twentymile Loop Trail in the Great Smoky Mountain National Park. Twelve soil specimens were collected from twelve different sites (**Figure 1A**) along the trail. A Scientific Research and Collecting Permit (Permit ID: GRSM-2021-SCI-2152) issued by the United Stated Department of the Interior National Park Service to conduct field research in GSMNP was obtained prior to sample collection and guidelines placed by the park were followed: collection from 15% or greater slopes and collection of less than 200g of organic soil per site. Samples were collected with sterile scoops and placed into sterile bags (Whirl-Pak, Madison, WI) while wearing sanitized gloves. Sample bags were kept in soft coolers with ice packs until arrival at the lab (<4 h), where they were then stored at 4°C until processing. For each sample collection, site location coordinates, terrain description, time, and sample mass were documented.

### *Listeria* spp. enrichment and isolation

Methods used to isolate *Listeria* spp. from the soil samples were adapted from the FDA Bacteriological Analytical Manual (BAM) (47). For each sample, 25 g of soil was added to 225 mL of Buffered Listeria Enrichment Broth (BLEB) (BD Difco, Franklin Lakes, New Jersey) and stomached at 260 RPM for one minute. The mixtures were then statically incubated at 30°C. After 4 h incubation, 900 μL of *Listeria* selective enrichment supplement (LSES, Oxoid, Thermo Fisher Scientific, Waltham, MA) was added to each enrichment and incubation was continued at 30°C. At 24 and 48 h of total incubation time, 1 mL aliquots of the BLEB enrichments were serially diluted to 10^-3^ in phosphate buffered saline (PBS; 1M; potassium chloride [Fischer Chemical], potassium phosphate [Acros Organics], sodium phosphate [Acros Organics], sodium chloride [Fischer Chemical], pH 7.4 with 6M HCl). 100 μL of each dilution was spread plated on Modified Oxford Agar (MOX; Oxford medium, Remel, Thermo Fisher Scientific, Waltham, MA; modified oxford antimicrobic supplement, BD Difco, Franklin Lakes, New Jersey) and incubated at 35°C for 24 h. A single colony characteristic of *Listeria* spp. was plucked from each MOX plate and added to 5 mL of Brain Heart Infusion (BHI; BD Bacto, Franklin Lakes, New Jersey) broth and incubated at 25°C in a shaking water bath for 24 h. After incubation, freezer stocks were prepared by adding 1 mL of overnight culture to 1 mL of BHI broth with 30% v/v glycerol; these were stored at −80°C.

### DNA extraction, sequencing, and genome assembly

DNA from the isolates was extracted using a Qiagen QIAamp DNA mini kit (Hilden, Germany) per the manufacturer’s protocol, with slight modifications such as including an RNase treatment (48). DNA concentration and quality were measured using a NanoDrop spectrophotometer (Thermo Fisher Scientific, Waltham, MA) and Qubit (Invitrogen, Thermo Fisher Scientific, Waltham, MA). Library preparation and sequencing were performed by the Microbial Genome Sequencing Center (MiGS; Pittsburgh, PA). Sequencing libraries were prepared using Nextera XT kits (Illumina, San Diego, CA). Sequencing was performed with the NextSeq 2000 platform, with 151 bp paired-end read chemistry. Raw reads were trimmed using Trimmomatic (v0.35; with the following parameters: ILLUMINACLIP:NexteraPE-PE.fa:2:30:10 LEADING:3 TRAILING:3 SLIDINGWINDOW:4:15 MINLEN:36) (49) and quality parameters assessed using FastQC (v0.11.7) (50). Trimmed reads were used to create genome assemblies with SPAdes (v3.12.0) (51). Assembly statistics were assessed using QUAST (v4.6.3) (52), BBMap (v38.08) (53), and SAMtools (v0.1.8) (54). Genome assemblies were checked to ensure that lengths (2.8-3.7 Mb) and G+C content (34-45%) were consistent with those expected for *Listeria* spp. (55, 56) and sufficient quality (number of contigs <100 and read coverage >50 ×).

### Species identification and characterization

For species-level identification, PYANI (v0.2.11) (57) was used to calculate average nucleotide identity between isolates and type strains (downloaded from NCBI) and bactaxR (v 0.1.0) (58) was used to create an ANI dendrogram. *Brochothrix* spp. were also included as an outgroup. Isolate pairs with a high ANI (>99.99%) were further evaluated using the CFSAN SNP Pipeline (v 1.0.1) (59). Species-level identifications were also evaluated using the ribosomal multilocus sequence typing (rMLST) tool available from PubMLST (60, 61) and the Type Strain Genome Server (TYGS) (62). MLST sequence type (ST) and clonal complex (CC), lineage, and serogroup were determined using the Listeria Sequence Typing tool available from Institut Pasteur (63-65).

## Data availability

Raw sequencing reads were submitted to the Sequencing Read Archive (SRA) and genome assemblies to GenBank on NCBI under BioProject PRJNA760531. BioSample IDs are listed in **Table 1**.

## ACKNOWLEDGMENTS

This research was supported by the University of Tennessee’s Office of Undergraduate Research Summer Undergraduate Research Internship Program. We would also like to acknowledge Geographic Information Systems (GIS) Specialist Eric Arnold with the University of Tennessee Knoxville Libraries for his help with creating the maps.

